# psntools - a Python package for protein structure network analysis

**DOI:** 10.1101/2022.02.07.479254

**Authors:** Valentina Sora, Matteo Tiberti, Elena Papaleo

## Abstract

The application of network theory to investigate protein structures and conformational ensembles through Protein Structure Networks (PSNs) has proven particularly insightful to study protein dynamics, the potentially disruptive effects of disease-related mutations, and allosteric mechanisms. Here, we present *psntools*, a novel Python package for downstream analysis of PSNs. *psntools* is completely PSN-agnostic, in contrast with several available tools in the community. *psntools* relies only on a few Python dependencies, most notably MDAnalysis and NetworkX, works without external software, and can be incorporated into Python-based analysis pipelines. We also present an example of the usage of *psntools* on a case of study of biological interest, which helped produce novel insights on the structural details of the interaction between BCL-xL and the BH3 motif of BECLIN-1. The *psntools* package and the data associated with the case study are available at https://github.com/ELELAB/psntools.

## Introduction

Dynamic conformational ensembles better describe proteins than static structures. ^1^ Proteins undergo several conformational changes, and their structural propensities can be affected by ‘perturbations’, such as point mutations and post-translational modifications (PTMs), as well as by the interaction with other biomolecules. Some perturbations can be propagated to sites far from the site where the perturbation occurred, often at the base of allostery^2–7^.

In this context, the study of protein structures or structural ensembles using concepts from the mathematical field of graph theory has proven helpful in understanding the fine-grained networks of interactions at play, the mechanisms of protein folding, and the determinants of allosteric phenomena ^8–11^. This field of application is broadly known as Protein Structure Networks (PSNs), namely graphs representing three-dimensional protein conformations through residue-residue pairwise interactions. Indeed, in a PSN, nodes identify residues, and edges are often described as physical interactions between them (such as hydrophobic interactions, salt bridges, or hydrogen bonds) or obey custom definitions of residue-residue contacts ^12,13^.

Furthermore, the application of PSNs to protein structural ensembles has proven insightful to investigate functional residues and conformational changes ^14–18^. This effectiveness has translated into an expansion of the arsenal of tools available to build protein structure networks (PSNs) according to different methodologies. However, most of these approaches are integrated into software devoted to studying conformational ensembles ^19–21^. Moreover, standalone, PSN-oriented suites usually include analyses tailored to the type of PSN that they implement ^13,22–25^. Finally, some available tools have a closed-source code ^24^ Furthermore, others do not rely on version-control systems for their open-source codebase ^22,25^. For this reason, we developed *psntools*, a Python package entirely dedicated to PSN analyses and able to handle virtually any kind of PSN. *psntools* has already proven helpful in one of its earlier versions in investigating the structural determinants of an allosteric mechanism involving the binding of PUMA to BCL-xL ^26^

### Salient features

*psntools* is entirely agnostic regarding the type of PSN provided, as long as the PSN represents a network of interactions among residues in a protein system.

Specifically, *psntools* uses data structures provided in the widely used Python packages MDAnalysis ^27,28^ and NetworkX ^29^ to construct a new hybrid object representing a PSN (implemented in the *PSN* class). In detail, MDAnalysis’ *Residue* instances are employed to describe nodes, while NetworkX’s *Graph* class is used to implement the network itself. This choice allows *psntools* to rely on two actively developed and thoroughly tested packages for its core objects. *psntools* then includes several functions to perform network analysis, summarize the results in data frames, write them out and generate visual reports to ease their interpretation (Figure 1).

**Figure 1.**
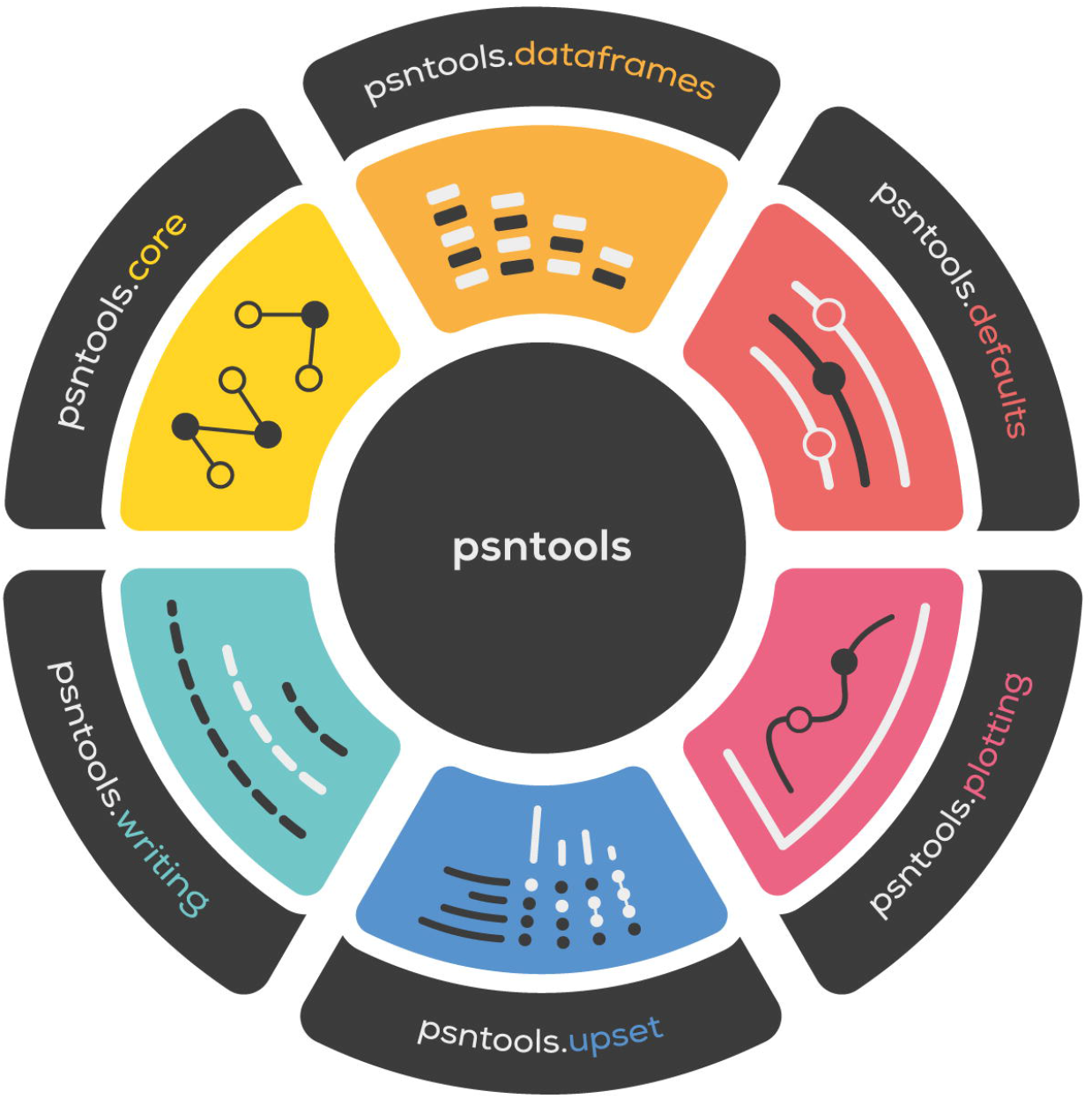
*psntools* modules available in the public API. The *psntools.defaults* module contains default hard-coded settings such as the path to the default directory where the configuration files for plotting can be found.

For instance, *psntools* can find hubs, connected components, shortest communication paths, and compute several centrality measures on single PSNs. Core objects and analysis functions all live in the *psntools.core* module. Results from all analyses can be either further manipulated as pure Python objects or converted into Pandas data frames using the functions provided in the *psntools.dataframes* module. Such data frames can then be formatted and written out to CSV files using the utilities in the *psntools.writing* module. A *psntools.plotting* module is also available, providing essential visualization utilities. Virtually all plot aesthetics can be tweaked via YAML configuration files passed directly to the plot functions. Examples of these configurations for different plot types are available inside the package in the *config_plot* folder.

Notably, *psntools* also includes *psntools.upset*, a module dedicated to an independent implementation of UpSet plots ^30^. UpSet plots are a particularly informative visualization strategy for set intersections. In *psntools*, these plots are employed when comparing several PSNs, easing the display of the nodes and edges that any possible combination of the given PSNs may have in common.

Indeed, *psntools* allows the representation of multiple PSNs as a *PSNGroup (PSNGroup* class). Gathering PSNs into a *PSNGroup* is most useful when the networks represent related systems. For example, they can be multiple conformational ensembles of the same system, a wild-type protein with several point mutants, or homologous proteins from different species. Indeed, a *PSNGroup* includes a one-to-one mapping of the nodes of a PSN to the nodes of all others included in the group. Therefore, nodes labeled as equal in different PSNs will be treated as they were the same when computing, for example, the number of nodes or edges common to a subset of PSNs in the group (or to all PSNs in the group). The concept of *PSNGroup* was introduced precisely to ease comparisons among PSNs where at least some nodes are equivalent to one another. This correspondence occurs, for instance, when nodes represent (i) the same residue or (ii) the same position (i.e., wild-type and mutated or otherwise modified) or (iii) equivalent sites (i.e., in homologous proteins). *PSNGroup* objects can be used to assess the changes in network properties upon mutation or in sets of related proteins, providing powerful functional insights about the systems of interest.

## Results

### Case study: structural determinants of the interaction between BCL-xL and the BH3 motif of BECLIN1

As mentioned in the Introduction, we first used *psntools* to investigate the structural details of an allosteric phenomenon triggered by the interaction between BCL-xL and PUMA ^26^, two proteins critical in the context of apoptosis, one of the mechanisms of programmed cell death^31^. In detail, BCL-xL belongs to the BCL-2 protein family and acts as a suppressor of apoptosis ^32^. On the other hand, PUMA belongs to the so-called BH3-only proteins, a group whose members interact with BCL-2 proteins through a short motif (the BH3 motif) and prevent them from performing their anti-apoptotic function ^33^.

Here, we present the features of *psntools* using as a case of study the interaction between BCL-xL and another BH3-only protein, BECLIN-1 ^34-36^ (Figure 3). This interaction is of particular interest not only in the context of apoptosis but also because it may constitute a contact point between apoptosis and autophagy, the cell’s primary recycling mechanism ^34,35,37^.

We decided to focus on comparing salt bridges, hydrogen bonds, and atomic-contacts-based PSNs found in the ensembles of both BCL-xL in its free state and of the complex between BCL-xL and the BH3 motif of BECLIN-1. We obtained these ensembles through one-μs Molecular Dynamics (MD) simulations using the CHARMM22* force field ^38^, and prepared as described in details for the MD of the BCL-xL:PUMA complex ^26^. The first model of the NMR structures with PDB IDs 2LPC ^39^ and 2PON ^34^ was used as a starting structure for the simulation of BCL-xL free and the complex, respectively. 2LPC covers residues 1-44 and 85-209 of BCL-xL (Figure 2, panel A), while 2PON covers residues 1-44 and 85-196 of BCL-xL, and residues 106-128 (the BH3 motif) of BECLIN-1 (Figure 2, panel B). A total of four ensembles (two for BCL-xL free, two for the complex) were used for PSN analyses. One of the ensembles of BCL-xL free was previously used in our study of the BCL-xL/PUMA interaction ^26^. It is referred to as “free_cap_tip3p” in the data associated with our previous publication (https://osf.io/rnc6d) and as “BCL-xL (2)” here. The other ensemble of BCL-xL (“BCL-xL (1)”) and the ensembles of the BCL-xL/BECLIN-1 complex (“BCL-xL/BECLIN-1 (1)” and “BCL-xL/BECLIN-1 (2)”) are presented here for the first time and have been included in the same OSF repository https://osf.io/rnc6d.

**Figure 2.**
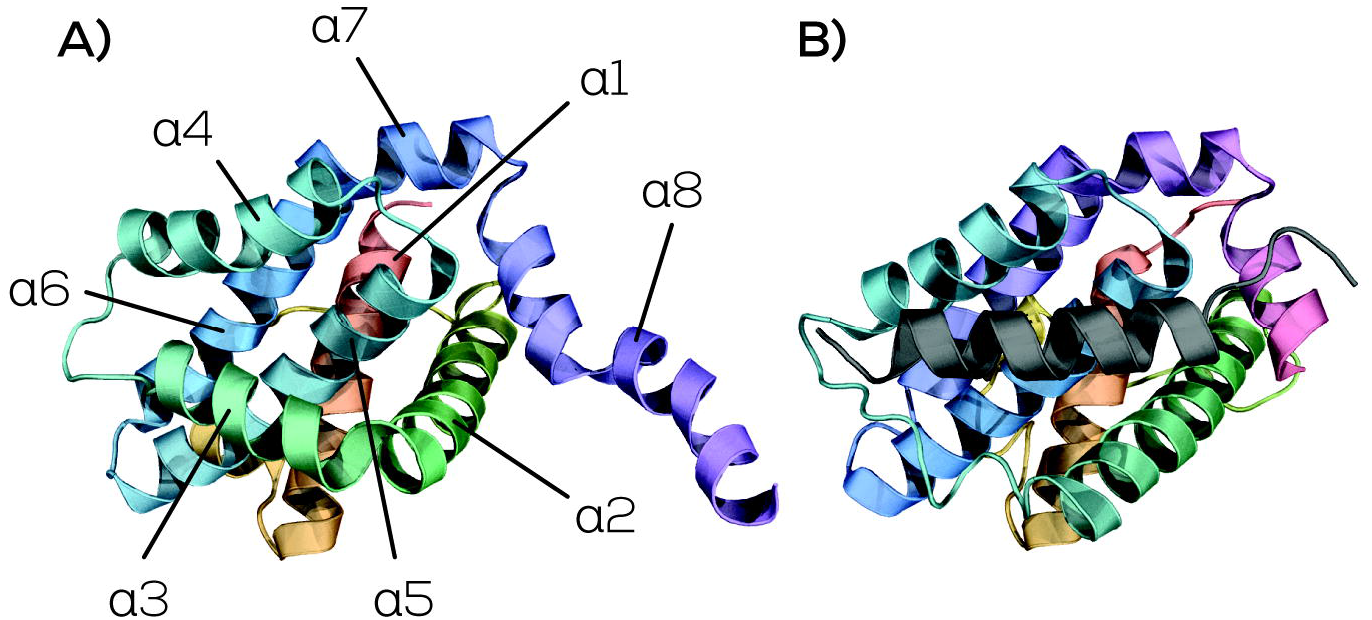
In panel A, the structure of Bcl-xL free (PDB ID: 2LPC) with the eight helices numbered. In panel B, the Bcl-xL/Beclin-1 complex (PDB ID: 2PON) with Bcl-xL shown in rainbow colors and Beclin-1 shown in dark gray.

### PSN calculation

Protein structure networks representing salt-bridged, hydrogen-bonded, and atomic-contacts-based interactions were calculated using PyInteraph2 ^40^. Only atoms belonging to charged moieties of two oppositely charged residues were considered for salt bridges calculation. We identified a salt bridge between two charged groups if we found at least one pair of atoms at a distance shorter than 5 Å. For aspartate and glutamate, the charged group was the carboxylic group. For lysine and arginine, the NH3- and the guanidinium groups were used, respectively. For a hydrogen bond to form, the distance between the acceptor atom and the hydrogen atom had to be lower than 3.5 Å, and the donor-hydrogen-acceptor atom angle greater than 120°. We calculated networks of main chain-main chain, main chain-side chain, and side chain-side chain hydrogen bonds.

In each PSN, each edge was weighted on the “occurrence” of the residue-residue contact it represented, defined as the percentage of structures in the ensemble (i.e., percentage of frames in the MD simulation) in which the contact was present. Only edges with a minimum occurrence of 20 % were kept in the final PSNs. This threshold was found to represent a good choice for retaining relevant interactions without including excessive noise ^13^. The atomic-contacts-based PSN (acPSN) is calculated by PyInteraph2 using an approach initially developed by Kannan and Vishveshwara ^12^, where the weight of each edge is computed as the number of atom pairs found in contacts between two residues, divided for the square root of the product of two “normalization factors”, one for each residue. Normalization factors for canonical residue types were presented in the original acPSN publication ^12^. Their relevance resides in the fact that they account for the residues’ propensities to form contacts. An occurrence cut-off of 50 % was used for acPSNs, the default used by the first implementation of acPSNs for conformational ensembles ^41^. The edge weight, called “interaction strength”, should exceed a pre-defined threshold for the edge to be reported in the final PSN. We followed the method proposed by Kannan and Vishveshwara to calculate the optimal interaction strength cut-off for each ensemble ^12^, which includes probing a range of values and calculating the size of the most populated component of the resulting PSNs. Connected components are isolated portions of the networks. Each node inside a connected component is connected (directly or indirectly) to all other nodes in its component and no other node outside the component. The size of the most populated of such components as a function of the interaction strength threshold should follow a sigmoidal behavior and abruptly drop around some value of interaction strength, identifying the point where the network sparsifies. We probed a range of values from 0.0 to 5.0 every 0.1 and computed the connected components using the *get_connected_components*() method of the PSN class in *psntools*. We found multiple drops occurring in this interval (Figure 3, panel A). However, the first significant reduction in size occurs around 2.4 and 2.7 for the PSNs of BCL-xL/BECLIN-1, while it happens around 3.1 for the BCL-xL free acPSNs. Therefore, we used these values as interaction strength cut-offs for our acPSNs.

**Figure 3.**
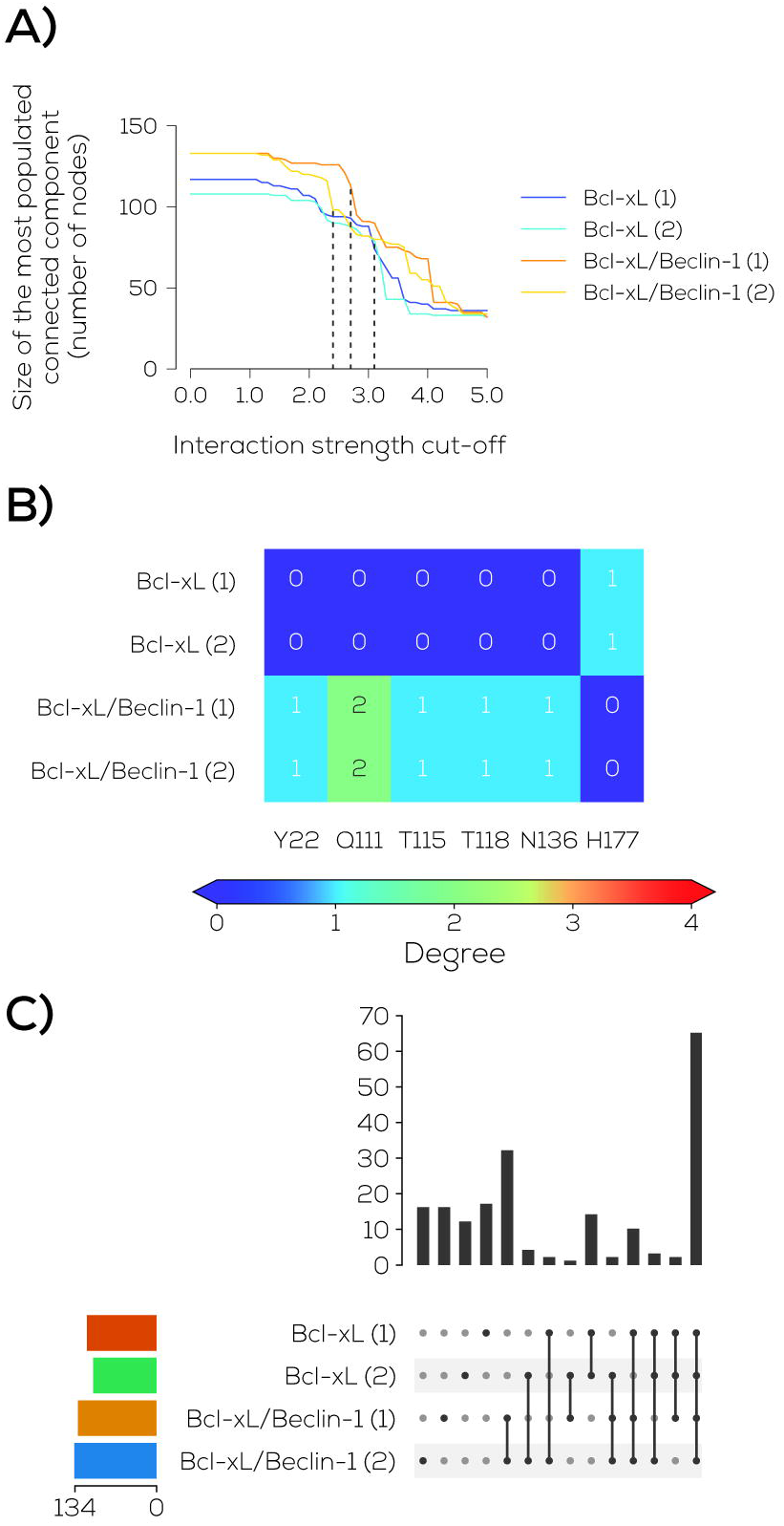
In panel A, the size of the most populated connected component at different interaction strength thresholds for acPSNs. In panel B, a heatmap highlighting the change in persistence values of the side chain - side chain hydrogen bonds across the different ensembles for nodes in Bcl-xL showing a rewiring of their network of hydrogen-bonded interactions. In panel C, an UpSet representation of the edges common to different subsets of acPSNs. The total number of edges for each acPSN is displayed by a colored horizontal bar on the left, while the sizes of the intersections between sets of edges in the different acPSNs are represented by the vertical bars on the right.

### Analyses with *psntools*

First, we generated a data frame containing edges that were common to each subset of PSNs for each type of network using the *psntools.dataframes.get_common_edges_df* function. We could therefore notice two highly persistent inter-chain salt bridges (> 94%) and side-chain – side-chain hydrogen bonds (> 80%) formed by K117 in BECLIN-1 with D133 and E129 in BCL-xL in both ensembles of the BCL-xL/BECLIN-1 complex. K117 is found to strongly interact with E129 and D133 in acPSNs (interaction strength > 6.6 and > 11.3, respectively). The interaction between K117 and E189 may be further stabilized by a weak (23.2-23.9 % occurrence) main chain – side-chain hydrogen bond. The relevance of K117 in mediating the interaction between the BH3 motif of BECLIN-1 and BCL-xL has also been highlighted by Feng and coworkers ^42^. Indeed, they demonstrated that the K117Q mutation destabilized the binding between BCL-xL and BECLIN-1.

On the other hand, one of the conserved positions in BECLIN-1’s BH3 motif, D121, is engaged in highly persistent side-chain – side-chain hydrogen bonds with R139 and N136 in BCL-xL. Furthermore, acPSNs identified D121-R139 as a strong contact (interaction strength > 12). The interaction between R139 and the conserved aspartate in BECLIN-1 has been observed before ^35,36,42^ and has also been found in other BCL-xL/BH3-motif complexes ^43–46^.

Then, we investigated changes in the BH3-binding groove in BCL-xL caused by the interaction with BECLIN-1 by comparing the number of contacts made by each node in BCL-xL in the different ensembles (the node “degree”). These alterations can be visually inspected through the *psntools.plotting.plot_nodes_df* function, which generates a heatmap displaying the degree of each node in different PSNs of the same type (see plots deposited in GitHub). We saw that most salt bridges were stable in all four ensembles from these heatmaps. In contrast, the network of side-chain – side-chain hydrogen bonds partially rewired upon the interaction with BECLIN-1. Therefore, we used *psntools.plotting.plot_nodes_df* to produce a more compact figure showing only nodes that gained or lost hydrogen bonds in the BCL-xL/BECLIN-1 PSNs with respect to the BCL-xL free PSNs (Figure 3, panel B). Interestingly, only some are in the BH3-binding groove (Q111, N136), while others are outside the pocket (Y22, T115, T118, H177), suggesting multiple structural changes throughout the protein to accommodate the BH3 motif.

Furthermore, we generated UpSet plots to investigate the shared hubs, intra-chain, and interchain edges among subsets of PSNs of the same type built from each of the four ensembles. This analysis showed 19 intra-chain edges shared by the acPSNs of BCL-xL free but absent in either of the acPSNs of the complex (Figure 3, panel C). In detail, six of these edges (L112-V126, H113-T115, L108-V126, Q111-V126, L105-F146, L108-F146) involve residues belonging to helix α3 of BCL-xL (first residue of each pair), which is highly flexible upon interaction with BH3 motifs ^35,44,47–50^. The loss of these edges in the acPSNs of the complex because they were either too weak or too transient further underlines this dynamic behavior in simulated ensembles of BCL-xL/BECLIN-1. Furthermore, V126 and F146 have been shown to mediate interactions between BCL-xL and other BH3 motifs ^43,46,51,52^, suggesting that the dynamics of helix a3 may be relevant in accommodating the binding partner in the groove.

### Conclusions and future developments

*psntools* represents a unicum in the landscape of PSN-related suites since it combines autonomy from external software, minimal dependencies, and the possibility of being seamlessly integrated into external Python-based analysis pipelines. Furthermore, its analysis and novel visualization capabilities have proven helpful in gaining insights and finding patterns in PSN data that help investigate the structure and dynamics of protein systems.

*psntools*’ structure makes it especially suitable for future incorporation into the MDAnalysis package to analyze conformational ensembles from molecular simulations. *psntools* could function as a plug-in extension installed with the main suite if needed. Pursuing this direction would not betray the original purpose of maintaining *psntools* as autonomous as possible since MDAnalysis is already one of the few dependencies required to install *psntools*. Furthermore, the software would benefit from being used and supported by a lively community, where a larger pool of users could report bugs and request new features. In this context, *psntools* could also benefit from being incorporated into the PyInteraph2 software ^40^, bringing the at-a-glance visualization utilities and the capability of grouping PSNs to an actively developed suite that already integrates several methodologies for PSN construction and would gain further appeal in the community from an extension of its analysis toolkit.

## Acknowledgments

This project was supported by Carlsberg Foundation Distinguished Fellowship (CF18-0314), Danmarks Grundforskningsfond (DNRF125), Hartmanns Fond (R421-A33877), LEO Foundation (LF17006), and NovoNordisk Fonden in Bioscience and Basic Biomedicine (0065262). The simulations were collected thanks to a DECI-PRACE 15^th^ HPC Grant on Archer.

## References

(1) Henzler-Wildman, K.; Kern, D. Dynamic Personalities of Proteins. Nature 2007, 450 (7172), 964–972. https://doi.org/10.1038/NATURE06522.

(2) Wodak, S. J.; Paci, E.; Dokholyan, N. v.; Berezovsky, I. N.; Horovitz, A.; Li, J.; Hilser, V. J.; Bahar, I.; Karanicolas, J.; Stock, G.; Hamm, P.; Stote, R. H.; Eberhardt, J.; Chebaro, Y.; Dejaegere, A.; Cecchini, M.; Changeux, J. P.; Bolhuis, P. G.; Vreede, J.; Faccioli, P.; Orioli, S.; Ravasio, R.; Yan, L.; Brito, C.; Wyart, M.; Gkeka, P.; Rivalta, I.; Palermo, G.; McCammon, J. A.; Panecka-Hofman, J.; Wade, R. C.; di Pizio, A.; Niv, M. Y.; Nussinov, R.; Tsai, C. J.; Jang, H.; Padhorny, D.; Kozakov, D.; McLeish, T. Allostery in Its Many Disguises: From Theory to Applications. Structure 2019, 27 (4), 566–578. https://doi.org/10.1016/j.str.2019.01.003.

(3) Papaleo, E.; Saladino, G.; Lambrughi, M.; Lindorff-Larsen, K.; Gervasio, F. L.; Nussinov, R. The Role of Protein Loops and Linkers in Conformational Dynamics and Allostery. Chemical reviews 2016, 116 (11), 6391–6423. https://doi.org/10.1021/acs.chemrev.5b00623.

(4) Abrusán, G.; Marsh, J. A. Ligand-Binding-Site Structure Shapes Allosteric Signal Transduction and the Evolution of Allostery in Protein Complexes. Molecular Biology and Evolution 2019, 36 (8), 1711–1727. https://doi.org/10.1093/molbev/msz093.

(5) Guarnera, E.; Berezovsky, I. N. On the Perturbation Nature of Allostery: Sites, Mutations, and Signal Modulation. Current opinion in structural biology 2019, 56, 18–27. https://doi.org/10.1016/J.SBI.2018.10.008.

(6) Naganathan, A. N. Modulation of Allosteric Coupling by Mutations: From Protein Dynamics and Packing to Altered Native Ensembles and Function. Current opinion in structural biology 2019, 54, 1–9. https://doi.org/10.1016/J.SBI.2018.09.004.

(7) Nussinov, R.; Tsai, C.-J. Allostery without a Conformational Change? Revisiting the Paradigm. Current Opinion in Structural Biology 2015, 30, 17–24. https://doi.org/10.1016/j.sbi.2014.11.005.

(8) di Paola, L.; Giuliani, A. Protein Contact Network Topology: A Natural Language for Allostery. Current Opinion in Structural Biology 2015, 31, 43–48. https://doi.org/10.1016/j.sbi.2015.03.001.

(9) Vishveshwara, S.; Brinda, K. V.; Kannan, N. Protein Structure: Insights From Graph Theory. Journal of Theoretical and Computational Chemistry 2002, 01 (01), 187–211. https://doi.org/10.1142/s0219633602000117.

(10) Vishveshwara, S.; Ghosh, A.; Hansia, P. Intra and Inter-Molecular Communications through Protein Structure Network. Current protein & peptide science 2009, 10 (2), 146–160. https://doi.org/10.2174/138920309787847590.

(11) Papaleo, E. Integrating Atomistic Molecular Dynamics Simulations, Experiments, and Network Analysis to Study Protein Dynamics: Strength in Unity. Frontiers in Molecular Biosciences 2015, 2 (May), 1–6. https://doi.org/10.3389/fmolb.2015.00028.

(12) Kannan, N.; Vishveshwara, S. Identification of Side-Chain Clusters in Protein Structures by a Graph Spectral Method. Journal of molecular biology 1999, 292 (2), 441–464. https://doi.org/10.1006/JMBI.1999.3058.

(13) Tiberti, M.; Invernizzi, G.; Lambrughi, M.; Inbar, Y.; Schreiber, G.; Papaleo, E. PyInteraph: A Framework for the Analysis of Interaction Networks in Structural Ensembles of Proteins. Journal of chemical information and modeling 2014, 54 (5), 1537–1551. https://doi.org/10.1021/CI400639R.

(14) Atilgan, A. R.; Akan, P.; Baysal, C. Small-World Communication of Residues and Significance for Protein Dynamics. Biophysical journal 2004, 86 (1 Pt 1), 85–91. https://doi.org/10.1016/S0006-3495(04)74086-2.

(15) Ghosh, A.; Vishveshwara, S. A Study of Communication Pathways in Methionyl-TRNA Synthetase by Molecular Dynamics Simulations and Structure Network Analysis. Proceedings of the National Academy of Sciences of the United States of America 2007, 104 (40), 15711–15716. https://doi.org/10.1073/PNAS.0704459104.

(16) Angelova, K.; Felline, A.; Lee, M.; Patel, M.; Puett, D.; Fanelli, F. Conserved Amino Acids Participate in the Structure Networks Deputed to Intramolecular Communication in the Lutropin Receptor. Cellular and molecular life sciences: CMLS 2011, 68 (7), 1227–1239. https://doi.org/10.1007/s00018-010-0519-z.

(17) Astl, L.; Verkhivker, G. M. Data-Driven Computational Analysis of Allosteric Proteins by Exploring Protein Dynamics, Residue Coevolution and Residue Interaction Networks. BBA - Gen Subj 2019, S0304-4165, 30179–5. https://doi.org/10.1016/j.bbagen.2019.07.008.

(18) Borsatto, A.; Marino, V.; Abrusci, G.; Lattanzi, G.; Dell’orco, D. Effects of Membrane and Biological Target on the Structural and Allosteric Properties of Recoverin: A Computational Approach. International journal of molecular sciences 2019, 20 (20). https://doi.org/10.3390/IJMS20205009.

(19) Brown, D. K.; Penkler, D. L.; Amamuddy, O. S.; Ross, C.; Atilgan, A. R.; Atilgan, C.; Bishop, Ö. T. MD-TASK: A Software Suite for Analyzing Molecular Dynamics Trajectories. Bioinformatics (Oxford, England) 2017, 33 (17), 2768–2771. https://doi.org/10.1093/BIOINFORMATICS/BTX349.

(20) Serçinoğlu, O.; Ozbek, P. GRINN: A Tool for Calculation of Residue Interaction Energies and Protein Energy Network Analysis of Molecular Dynamics Simulations. Nucleic acids research 2018, 46 (W1), W554–W562. https://doi.org/10.1093/NAR/GKY381.

(21) Grant, B. J.; Skjærven, L.; Yao, X. Q. The Bio3D Packages for Structural Bioinformatics. Protein science: a publication of the Protein Society 2021, 30 (1), 20–30. https://doi.org/10.1002/PRO.3923.

(22) Bhattacharyya, M.; Bhat, C. R.; Vishveshwara, S. An Automated Approach to Network Features of Protein Structure Ensembles. Protein sciencexg: a publication of the Protein Society 2013, 22 (10), 1399–1416. https://doi.org/10.1002/PRO.2333.

(23) Contreras-Riquelme, S.; Garate, J. A.; Perez-Acle, T.; Martin, A. J. M. RIP-MD: A Tool to Study Residue Interaction Networks in Protein Molecular Dynamics. PeerJ 2018, 6 (12). https://doi.org/10.7717/PEERJ.5998.

(24) Chakrabarty, B.; Parekh, N. NAPS: Network Analysis of Protein Structures. Nucleic Acids Research 2016, 44 (W1), W375–W382. https://doi.org/10.1093/nar/gkw383.

(25) Felline, A.; Seeber, M.; Fanelli, F. PSNtools for Standalone and Web-Based Structure Network Analyses of Conformational Ensembles. Computational and Structural Biotechnology Journal 2022, 20, 640–649. https://doi.org/10.1016/j.csbj.2021.12.044.

(26) Sora, V.; Sanchez, D.; Papaleo, E. Bcl-XL Dynamics under the Lens of Protein Structure Networks. The journal of physical chemistry. B 2021, 125 (17), 4308–4320. https://doi.org/10.1021/ACS.JPCB.0C11562.

(27) Michaud-Agrawal, N.; Denning, E. J.; Woolf, T. B.; Beckstein, O. MDAnalysis: A Toolkit for the Analysis of Molecular Dynamics Simulations. Journal of computational chemistry 2011, 32 (10), 2319–2327. https://doi.org/10.1002/JCC.21787.

(28) Gowers, R.; Linke, M.; Barnoud, J.; Reddy, T.; Melo, M.; Seyler, S.; Domański, J.; Dotson, D.; Buchoux, S.; Kenney, I.; Beckstein, O. MDAnalysis: A Python Package for the Rapid Analysis of Molecular Dynamics Simulations. Proceedings of the 15th Python in Science Conference 2016, No. Scipy, 98–105. https://doi.org/10.25080/majora-629e541a-00e.

(29) Hagberg, A. A.; Schult, D. A.; Swart, P. J. Exploring Network Structure, Dynamics, and Function Using NetworkX. 7th Python in Science Conference (SciPy 2008) 2008, No. SciPy, 11–15.

(30) Lex, A.; Gehlenborg, N.; Strobelt, H. UpSet: Visualization of Intersecting Sets. IEEE Trans Vis Comput Graph. 2014, 20 (12), 1983–1992. https://doi.org/10.1109/TVCG.2014.2346248.UpSet.

(31) Czabotar, P. E.; Lessene, G.; Strasser, A.; Adams, J. M. Control of Apoptosis by the BCL-2 Protein Family: Implications for Physiology and Therapy. Nature reviews. Molecular cell biology 2014, 15 (1), 49–63. https://doi.org/10.1038/nrm3722.

(32) Singh, R.; Letai, A.; Sarosiek, K. Regulation of Apoptosis in Health and Disease: The Balancing Act of BCL-2 Family Proteins. Nature reviews. Molecular cell biology 2019, 20 (3), 175–193. https://doi.org/10.1038/S41580-018-0089-8.

(33) Huang, K.; O’Neill, K. L.; Li, J.; Zhou, W.; Han, N.; Pang, X.; Wu, W.; Struble, L.; Borgstahl, G.; Liu, Z.; Zhang, L.; Luo, X. BH3-Only Proteins Target BCL-XL/MCL-1, Not BAX/BAK, to Initiate Apoptosis. Cell research 2019, 29 (11), 942–952. https://doi.org/10.1038/S41422-019-0231-Y.

(34) Feng, W.; Huang, S.; Wu, H.; Zhang, M. Molecular Basis of Bcl-XL’s Target Recognition Versatility Revealed by the Structure of Bcl-XL in Complex with the BH3 Domain of Beclin-1. Journal of molecular biology 2007, 372 (1), 223–235. https://doi.org/10.1016/J.JMB.2007.06.069.

(35) Oberstein, A.; Jeffrey, P. D.; Shi, Y. Crystal Structure of the Bcl-XL-Beclin 1 Peptide Complex: Beclin 1 Is a Novel BH3-Only Protein. The Journal of biological chemistry 2007, 282 (17), 13123–13132. https://doi.org/10.1074/JBC.M700492200.

(36) Lee, E. F.; Smith, N. A.; Soares da Costa, T. P.; Meftahi, N.; Yao, S.; Harris, T. J.; Tran, S.; Pettikiriarachchi, A.; Perugini, M. A.; Keizer, D. W.; Evangelista, M.; Smith, B. J.; Fairlie, W. D. Structural Insights into BCL2 Pro-Survival Protein Interactions with the Key Autophagy Regulator BECN1 Following Phosphorylation by STK4/MST1. Autophagy 2019, 15 (5), 785–795. https://doi.org/10.1080/15548627.2018.1564557.

(37) Maiuri, M. C.; Criollo, A.; Tasdemir, E.; Vicencio, J. M.; Tajeddine, N.; Hickman, J. A.; Geneste, O.; Kroemer, G. BH3-Only Proteins and BH3 Mimetics Induce Autophagy by Competitively Disrupting the Interaction between Beclin 1 and Bcl-2/Bcl-XL. https://doi.org/10.4161/auto.4237 2007, 3 (4), 374–376. https://doi.org/10.4161/AUTO.4237.

(38) Piana, S.; Lindorff-Larsen, K.; Shaw, D. E. How Robust Are Protein Folding Simulations with Respect to Force Field Parameterization? Biophysical journal 2011, 100 (9). https://doi.org/10.1016/J.BPJ.2011.03.051.

(39) Wysoczanski, P.; Mart, R. J.; Loveridge, E. J.; Williams, C.; Whittaker, S. B. M.; Crump, M. P.; Allemann, R. K. NMR Solution Structure of a Photoswitchable Apoptosis Activating Bak Peptide Bound to Bcl-x L. Journal of the American Chemical Society 2012, 134 (18), 7644–7647. https://doi.org/10.1021/JA302390A/SUPPL_FILE/JA302390A_SI_001.PDF.

(40) Sora, V.; Tiberti, M.; Mahdi Robbani, S.; Rubin, J.; Papaleo, E. PyInteraph2 and PyInKnife2 to Analyze Networks in Protein Structural Ensembles. bioRxiv 2020, 2020.11.22.381616.

(41) Seeber, M.; Felline, A.; Raimondi, F.; Muff, S.; Friedman, R.; Rao, F.; Caflisch, A.; Fanelli, F. Wordom: A User-Friendly Program for the Analysis of Molecular Structures, Trajectories, and Free Energy Surfaces. Journal of computational chemistry 2011, 32 (6), 1183–1194. https://doi.org/10.1002/JCC.21688.

(42) Feng, W.; Huang, S.; Wu, H.; Zhang, M. Molecular Basis of Bcl-XL’s Target Recognition Versatility Revealed by the Structure of Bcl-XL in Complex with the BH3 Domain of Beclin-1. Journal of Molecular Biology 2007, 372 (1), 223–235. https://doi.org/10.1016/j.jmb.2007.06.069.

(43) Sattler, M.; Liang, H.; Nettesheim, D.; Meadows, R. P.; Harlan, J. E.; Eberstadt, M.; Yoon, H. S.; Shuker, S. B.; Chang, B. S.; Minn, A. J.; Thompson, C. B.; Fesik, S. W. Structure of Bcl-XL-Bak Peptide Complex: Recognition between Regulators of Apoptosis. Science (New York, N.Y.) 1997, 275 (5302), 983–986. https://doi.org/10.1126/SCIENCE.275.5302.983.

(44) Boersma, M. D.; Haase, H. S.; Peterson-Kaufman, K. J.; Lee, E. F.; Clarke, O. B.; Colman, P. M.; Smith, B. J.; Horne, W. S.; Fairlie, W. D.; Gellman, S. H. Evaluation of Diverse α/β-Backbone Patterns for Functional α-Helix Mimicry: Analogues of the Bim BH3 Domain. Journal of the American Chemical Society 2012, 134 (1), 315–323. https://doi.org/10.1021/JA207148M.

(45) Rajan, S.; Choi, M.; Baek, K.; Yoon, H. S. Bh3 Induced Conformational Changes in Bcl-Xl Revealed by Crystal Structure and Comparative Analysis. Proteins 2015, 83 (7), 1262–1272. https://doi.org/10.1002/PROT.24816.

(46) Thebault, S.; Agez, M.; Chi, X.; Stojko, J.; Cura, V.; Telerman, S. B.; Maillet, L.; Gautier, F.; Billas-Massobrio, I.; Birck, C.; Troffer-Charlier, N.; Karafin, T.; Honore, J.; Senff-Ribeiro, A.; Montessuit, S.; Johnson, C. M.; Juin, P.; Cianferani, S.; Martinou, J. C.; Andrews, D. W.; Amson, R.; Telerman, A.; Cavarelli, J. TCTP Contains a BH3-like Domain, Which Instead of Inhibiting, Activates Bcl-XL. Scientific reports 2016, 6. https://doi.org/10.1038/SREP19725.

(47) Liu, X.; Dai, S.; Zhu, Y.; Marrack, P.; Kappler, J. W. The Structure of a Bcl-XL/Bim Fragment Complex: Implications for Bim Function. Immunity 2003, 19 (3), 341–352. https://doi.org/10.1016/S1074-7613(03)00234-6.

(48) Lee, E. F.; Fedorova, A.; Zobel, K.; Boyle, M. J.; Yang, H.; Perugini, M. A.; Colman, P. M.; Huang, D. C. S.; Deshayes, K.; Fairlie, W. D. Novel Bcl-2 Homology-3 Domain-like Sequences Identified from Screening Randomized Peptide Libraries for Inhibitors of the Pro-Survival Bcl-2 Proteins * □. 2009, 284 (45), 31315–31326. https://doi.org/10.1074/jbc.M109.048009.

(49) Follis, A. V.; Chipuk, J. E.; Fisher, J. C.; Yun, M. K.; Grace, C. R.; Nourse, A.; Baran, K.; Ou, L.; Min, L.; White, S. W.; Green, D. R.; Kriwacki, R. W. PUMA Binding Induces Partial Unfolding within BCL-XL to Disrupt P53 Binding and Promote Apoptosis. Nature chemical biology 2013, 9 (3), 163–168. https://doi.org/10.1038/NCHEMBIO.1166.

(50) Lee, D.-H.; Ha, J.-H.; Kim, Y.; Jang, M.; Park, S. J.; Yoon, H. S.; Kim, E.-H.; Bae, K.-H.; Park, B. C.; Park, S. G.; Yi, G.-S.; Chi, S.-W. A Conserved Mechanism for Binding of P53 DNA-Binding Domain and Anti-Apoptotic Bcl-2 Family Proteins. Molecules and cells 2014, 37 (3), 264–269. https://doi.org/10.14348/molcells.2014.0001.

(51) Petros, A. M.; Nettesheim, D. G.; Wang, Y.; Olejniczak, E. T.; Meadows, R. P.; Mack, J.; Swift, K.; Matayoshi, E. D.; Zhang, H.; Fesik, S. W.; Thompson, C. B. Rationale for Bcl-XL/Bad Peptide Complex Formation from Structure, Mutagenesis, and Biophysical Studies. Protein science: a publication of the Protein Society 2000, 9 (12), 2528–2534. https://doi.org/10.1110/PS.9.12.2528.

(52) Ambrosi, E.; Capaldi, S.; Bovi, M.; Saccomani, G.; Perduca, M. Structural Changes in the BH3 Domain of SOUL Protein upon Interaction with the Anti-Apoptotic Protein Bcl-XL. 2011, 301, 291–301. https://doi.org/10.1042/BJ20110257.

